# Broad brain biodistribution conferred by an AAV to restore TDP-43 function mitigates Frontotemporal Demenia-like deficits

**DOI:** 10.1101/2025.08.28.672900

**Authors:** Tianyu Cao, Rashmi Thapa, Rongsong Liu, Aswathy Peethambaran Mallika, Meghraj Singh Baghel, Yijun Wei, Irika R. Sinha, Grace D. Burns, Xinrui Wen, Bo Pang, Jonathan P Ling, Da-Ting Lin, Yun Li, Philip C. Wong

**Author notes:** **Corresponding author: P.C.W. (**).

## Abstract

TDP-43 dysfunction is an early pathogenic determinant of frontotemporal lobar degeneration with TDP-43 pathology (FTLD-TDP), a devastating disorder currently without effective therapy. Here, we exploit a blood-brain-barrier (BBB)-permeable AAV (AAV-PHP.eB) that confers broad brain biodistribution to restore TDP-43 function in a TDP-43 deficient model (*CamKIIa-CreER;Tardbp* mice) that mimics the early stage of TDP-43 dysfunction occurring in FTLD-TDP. Intracerebroventricular delivery by AAV-PHP.eB of CTR, our previously characterized splicing repressor, revealed its accumulation in ∼40% of adult hippocampal neurons. Remarkably, treatment of adult *CamKIIa-CreER;Tardbp^f/f^*mice with AAV-PHP.eB-CTR restored TDP-43 function, attenuated neuronal aberrant activity and memory deficits, and rescued neuron loss. Importantly, we showed that TDP-43’s autoregulatory element restricts CTR expression to a physiological range. No overt phenotype was observed after long-term exposure to AAV-PHP.eB-CTR in aged mice, highlighting a favorable safety profile for this gene therapy. These results validate that BBB-crossing AAVs can deliver CTR with a biodistribution in the adult brain that is broad enough to rescue FTD-like phenotypes, supporting clinical testing of this gene therapy for FTLD-TDP.

## Introduction

Emerging evidence^1–3^ supports the notion that loss of nuclear TAR DNA/RNA-binding protein 43kDa (*TARDBP*, TDP-43) and its splicing repression underlies frontotemporal dementia (FTD) and amyotrophic lateral sclerosis (ALS), devastating adult-onset neurodegenerative disorders^4–12^ currently without disease modifying therapy. TDP-43, a highly conserved nuclear RNA binding protein (RBP), was first proposed to primarily induce cytoplasmic neuronal inclusions and drive neuron loss through such aggregates in this disease spectrum^13^. A major neuronal function of TDP-43 was subsequently shown to be the regulation of cryptic exons splicing, the loss of which is implicated in ALS-FTD^1^. This discovery led to the proposal that deficits in TDP-43 cryptic splicing can be an early pathogenic event that drives neuron loss. This view is supported by observations that cryptic exons encoded peptides, such as that found in cryptic *HDGFL2*, can be found in biofluids of ALS-FTD patients during early and presymptomatic stages of disease, indicative of early loss of TDP-43 function^2,3^. Findings of TDP-43 splicing dysfunction preceding cytoplasmic inclusion in the human aging brain by at least a decade^14^ also supports this view. This model is further strengthened by the observations that mutations in TDP-43 linked to ALS^15–17^ impact a TDP-43 cryptic exon of stathmin-2 (*STMN2*) in human iPSC derived neurons independent of TDP-43 cytoplasmic aggregates^18–20^. *UNC13A*, another TDP-43 target, encodes a strong risk allele for ALS and FTLD-TDP which influences the inclusion of its cryptic exon^21,22^. Observations of neuronal TDP-43 nuclear depletion in a presymptomatic C9orf72 patient ^23^ and some FTLD-TDP cases^24^, along with cryptic exons in FTLD-TDP and AD-TDP cases^25,26^, all without cytoplasmic aggregates, provide additional data. These data support a model that deficits associated with TDP-43 cryptic splicing is triggered during the presymptomatic stage of disease to drive neuron loss. Thus, therapeutic strategies designed to restore TDP-43 function represent a potential mechanism-based therapy for FTLD-TDP.

Using our conditional knockout mouse model with TDP-43 depletion in forebrain neurons (*CamKIIa-CreER;Tardbp^f/f^*) that exhibits selective neuron loss mimicking the prodromal phase of FTLD-TDP, we previously showed that loss of TDP-43 dependent cryptic splicing leads to activation of caspase-3^27,28^, forebrain circuit abnormalities and memory deficits^29^. Importantly, we developed an AAV-based therapeutic strategy to restore the loss of splicing repression by TDP-43 using a novel chimeric splicing repressor, termed CTR^1,30^. To validate the therapeutic efficacy and potential toxicity associated with such AAV-mediated delivery of CTR to forebrain neurons, we delivered CTR intracerebroventricularly to adult *CamKIIa-CreER;Tardbp^f/f^* mice utilizing a BBB crossing serotype, AAV-PHP.eB^31,32^ that facilitates efficient transduction of adult central neurons. We report here the safety and efficacy of AAV-PHP.eB-CTR in restoring TDP-43 function and attenuating forebrain neuron loss and rescuing neuronal circuit and cognitive deficits. Importantly, the inclusion of TDP-43 autoregulatory element as a “safety switch” in the payload ensured CTR levels remained within the normal range in hippocampal neurons. Furthermore, long-term exposure of AAV-PHP.eB-CTR in aged mice showed no evidence of untoward events. These outcomes strongly support the favorable safety profile of this gene therapy. Together, this validation serves as a crucial step toward establishing the clinical viability of AAV-mediated CTR delivery as a potential therapy for FTLD-TDP.

## Results

### Intracerebroventricular delivery of AAV-PHP.eB-CTR broadly transduces adult forebrain neurons without excessive accumulation of CTR

We previously demonstrated that AAV9 delivery of CTR to P0 mice lacking TDP-43 in spinal motor neurons can complements TDP-43 loss of function (LOF)^30^. Build on this, the current study asked whether restoration of TDP-43’s splicing repression function could mitigate downstream consequences of TDP-43 loss in the *CamKIIa-CreER;Tardbp^f/f^* mouse model, which recapitulates early features ofFTLD-TDP, while also avoiding the known risks associated with TDP-43 overexpression and aggregation toxicity^33^. To achieve this, we designed a chimeric therapeutic construct, termed CTR^1^, which retains the N-terminal RNA recognition domains (RRM1 and RRM2, amino acids 1–267, ∼30 kDa) of TDP-43 to preserve its RNA-binding specificity, while replacing the glycine-rich C-terminal domain, which is implicated in pathological aggregation, with an unrelated splicing repressor domain from RAVER1^34–36^ (amino acids 450–643, ∼20 kDa), an unrelated RBP(**Supplementary figure 1C**).

Physiological regulation of expression was ensured by incorporating the 3′ untranslated region (3′UTR) of human *TARDBP*, which contains the polyadenylation-dependent autoregulatory element that enables TDP-43 to bind its own transcript and limit its expression levels^37,38^ (**Supplementary figure 1C**). This negative feedback mechanism is critical for maintaining normal nuclear TDP-43 homeostasis and preventing toxic accumulation and essential for neuronal survival^39^. Our RT-PCR analysis (**Fig. 2G**) confirmed that CTR expression is higher in forebrains of TDP-43–deficient (*CamKII-CreER;Tardbp^f/f^*) animals as compared to that of *Tardbp^f/f^* control littermates, indicating preserved autoregulation via this 3′UTR of *TARDBP*.

To validate such ability of CTR to rescue cell death of forebrain neurons lacking TDP-43^27,29^, we delivered intracerebroventricularly (ICV) AAV9-CTR to P0 *CamKIIa-CreER;Tardbp^f/f^* pups. We found that AAV9-CTR attenuates neuron loss occurring in CA2/3 hippocampal neurons of mice lacking TDP-43 (**Supplementary figure 2A-C**). Corroborating these results, we showed that TDP-43 cryptic exons are suppressed in rescued mice (**Supplementary figure 2D-F**), confirming that the failure to repress TDP-43 cryptic exons in central neurons underlies neuron loss. However, for clinically relevant context, it is critical to establish delivery of CTR using an AAV serotype that broadly transduces central neurons to restore TDP-43 LOF in the adult brain.

For efficient CTR delivery in adult forebrain neurons of *CamKIIa-CreER;Tardbp^f/f^* mice, we used a BBB crossing AAV-PHP.eB that facilitates efficient transduction of adult central neurons^31,32^ via the ICV route of administration. To model the early stage of TDP-43 LOF occurring in human disease^2^, we selectively deleted TDP-43 from forebrain neurons of adult *CamKIIa-CreER;Tardbp^f/f^*mice. Upon tamoxifen administration, Cre recombinase was activated in Camk2a-positive excitatory neurons, resulting in the excision of exon 3 in *Tardbp* and subsequent loss of TDP-43 in targeted regions of the brain (**Supplementary figure 1A-B**). Immunostaining confirmed robust nuclear depletion of TDP-43 in the hippocampus one month after tamoxifen exposure in 5 month-old *CamKIIa-CreER;Tardbp^f/f^* mice, whereas control littermates (*Tardbp^f/f^*mice) maintained normal level of TDP-43 (**Fig. 1A**).

**Figure 1.**
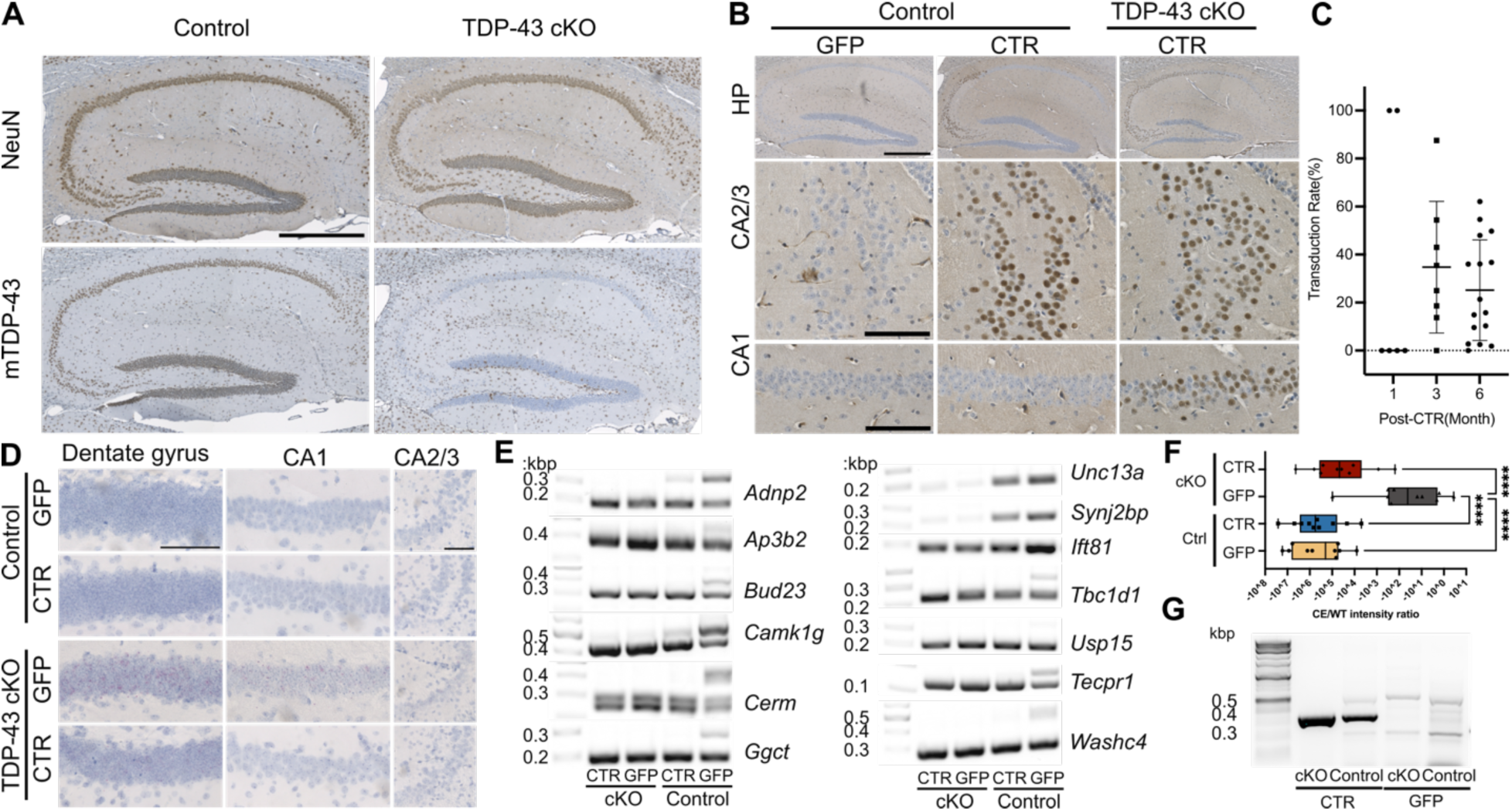
CTR is broadly expressed in neurons of adult brain and restores cryptic exon repression. (A) Representative immunohistochemistry images showing TDP-43 expression in the hippocampus 1 month after tamoxifen treatment. TDP-43 expression is retained in *Tardbp^f/f^* mice, but selectively lost in the TDP-43 conditional knockout (cKO) hippocampus, confirming efficient Cre recombination. Scale bar: 500 μm (B), Representative immunohistochemistry showing widespread CTR protein expression in the hippocampus of TDP-43 CKO and control mice following ICV of AAV-CTR. CTR expression was detected using an antibody targeting the human TDP-43 N-terminus. Strong expression was observed in the CA2 and CA3 subregions, with additional signal in CA1. (C), Quantification of CTR viral transduction rate in TDP-43 CKO mice at 1-, 3-, and 6-months post-injection(n = 6,8,16 respectively). Infection rates remained stable over time (∼40%). (D), Representative images of BaseScope *in situ* hybridization for cryptic Unc13a RNA reveals spatial distribution of splicing defects across hippocampal subregions. Cryptic Unc13a transcripts were elevated in dentate gyrus (DG), CA1, and CA2/3 of TDP-43 cKO-GFP mice and markedly reduced in CTR-treated animals, indicative of CTR restoring TDP-43 splicing regulation. (E), Representative RT-PCR analysis of hippocampal RNA showing repression of multiple cryptic exon–containing transcripts in CTR-treated TDP-43 CKO mice. Representative targets include *Adnp2, Ap3b2, Bud23, Camk1g, Cerm, Ggct, Unc13a, Synj2bp, Tbc1d1, Usp15, Tecpr1,* and *Washc4*. (F) Quantification of cryptic exon inclusion relative to wild-type transcript levels, expressed as the ratio of cryptic exon band intensity to corresponding wild-type band intensity from RT-PCR analyses (as in panel E). TDP-43 cKO mice treated with GFP exhibited a significant increase in cryptic exon usage compared to CTR-treated cKO and control groups, whereas CTR treated cKO mice were indistinguishable from control littermates. (G), RT-PCR quantification of CTR mRNA levels using primers spanning the TDP-43 N-terminal and RAVER1 C-terminal sequences. Despite equal AAV dosing, CTR transcript levels were significantly higher in TDP-43 cKO mice than in controls, consistent with autoregulatory stabilization of expression via the Tardbp 3′UTR. Scale bars: a: upper panel 500 μm, lower panel 100 μm; d: 100 μm. Error bars represent mean ± s.e.m. P-values were calculated using two-tailed unpaired t-tests or Mann–Whitney tests as specified in figure panels; n values are defined in the corresponding Methods.

### Delivery of AAV-PHP.eB-CTR to adult brain restores splicing repression in TDP-43 deficient hippocampal neurons

To evaluate the *in vivo* expression and function of the chimeric repressor CTR, we first assessed its distribution following ICV delivery of AAV-PHP.eB–CTR in *CamKIIa-CreER;Tardbp^f/f^* or *Tardbp^f/f^* mice. Immunohistochemical analysis using an antibody specific to the N-terminus of human TDP-43 revealed widespread expression of the CTR fusion protein throughout the hippocampus, with robust signal detected in CA2 and CA2/3 subregions and detectable expression in CA1 (**Fig. 1B**); quantification of transduction efficiency showed a stable transduction rate of approximately 40% of neurons in *CamKIIa-CreER;Tardbp^f/f^* mice (**Fig. 1C).**

We next asked whether CTR expression was sufficient to restore TDP-43’s splicing repressor function *in vivo*. RT-PCR of hippocampal tissue from treated *CamKIIa-CreER;Tardbp^f/f^* mice revealed a marked reduction in a panel of TDP-43–regulated cryptic exons, including those in *Adnp2, Ap3b2, Bud23, Camk1g, Cerm, Ggct, Unc13a, Synj2bp, Tbc1d1, Usp15, Tecpr1,* and *Washc4* (**Fig. 1E,F**), indicating broad repression of aberrant cryptic splicing. These cryptic exons span a range of genes involved in neuronal function, and are known to be directly regulated by TDP-43 in the mouse brain. Their coordinated repression by CTR suggests that the chimeric construct can functionally reconstitute TDP-43’s splicing repressor activity across multiple endogenous targets. This broad repression pattern reflects the molecular reach of CTR in restoring splicing fidelity in TDP-43–deficient CNS neurons when delivered to the adult brain.

To further validate cryptic exon repression at the cellular level, we performed *in situ* hybridization using a BaseScope probe targeting the cryptic exon in mouse *Unc13a*. *UNC13A* is a known ALS genetic risk factor and harbors a TDP-43–sensitive cryptic exons that are not conserved between mouse (within intron 1) and human (within intron 22)^22^. Upon TDP-43 depletion, cryptic *Unc13a* RNA accumulated in multiple hippocampal regions, with the highest signal in the dentate gyrus (DG), followed by CA1 and CA2/3 (**Fig. 1D**). This regional pattern may reflect differential vulnerability to TDP-43 depletion within the hippocampus. Notably, CTR treatment of *CamKIIa-CreER;Tardbp^f/f^* significantly reduced cryptic *Unc13a* expression in all subregions as compared to those of GFP-treated *CamKIIa-CreER;Tardbpf/f* mice (**Fig. 1D),** providing evidence that CTR effectively represses disease-relevant cryptic exons *in vivo*.

To ensure CTR expression is subject to TDP-43–like autoregulation, we analyzed CTR mRNA levels using RT-PCR with a forward primer targeting the TDP-43 N-terminus and a reverse primer targeting the RAVER1 C-terminus. Despite equivalent AAV doses, CTR transcript levels were significantly higher in *CamKIIa-CreER;Tardbp^f/f^*mice as compared to that of *Tardbp^f/f^* control littermates (**Fig. 1G**), consistent with the increased protein expression observed by immunohistochemical analysis (**Fig. 1B**). This result suggests that CTR retains the autoregulatory features of endogenous TDP-43 via its 3′UTR, leading to transcript stabilization and translation in the context of TDP-43 loss.

### AAV-PHP.eB-CTR ameliorates CA2/3 neuron loss and hippocampal atrophy

To assess the long-term neuroprotective effects of CTR in the context of TDP-43 loss, we examined neuronal survival in the hippocampus of *CamKIIa-CreER;Tardbp^f/f^* mice 12 months post-treatment. Immunohistochemical analysis using antibody recognizing NeuN revealed severe neuronal loss of ∼60% in the CA2/3 region of *CamKIIa-CreER;Tardbp^f/f^*mice treated with AAV-PHP.eB-GFP, with near-complete depletion of NeuN-positive cells compared to control *Tardbp^f/f^* mice (**Fig. 2A**). In contrast, ∼70% of neurons survive in CamKIIa-CreER;Tardbp^f/f^ mice treated with AAV-PHP.eB-CTR, (**Fig. 2C)**. Indicating that, on average, AAV-PHP.eB-CTR treated *CamKIIa-CreER;Tardbp^f/f^*mice protected ∼30% of the CA2/3 neurons from death, which closely matched the local transduction rate observed for AAV-PHP.eB-CTR (**Fig. 2C**). The extent of neuronal rescue in CA2/3 is consistent with the ∼40% transduction rate in this area, supporting a direct impact between CTR accumulation and cell survival.

**Figure 2.**
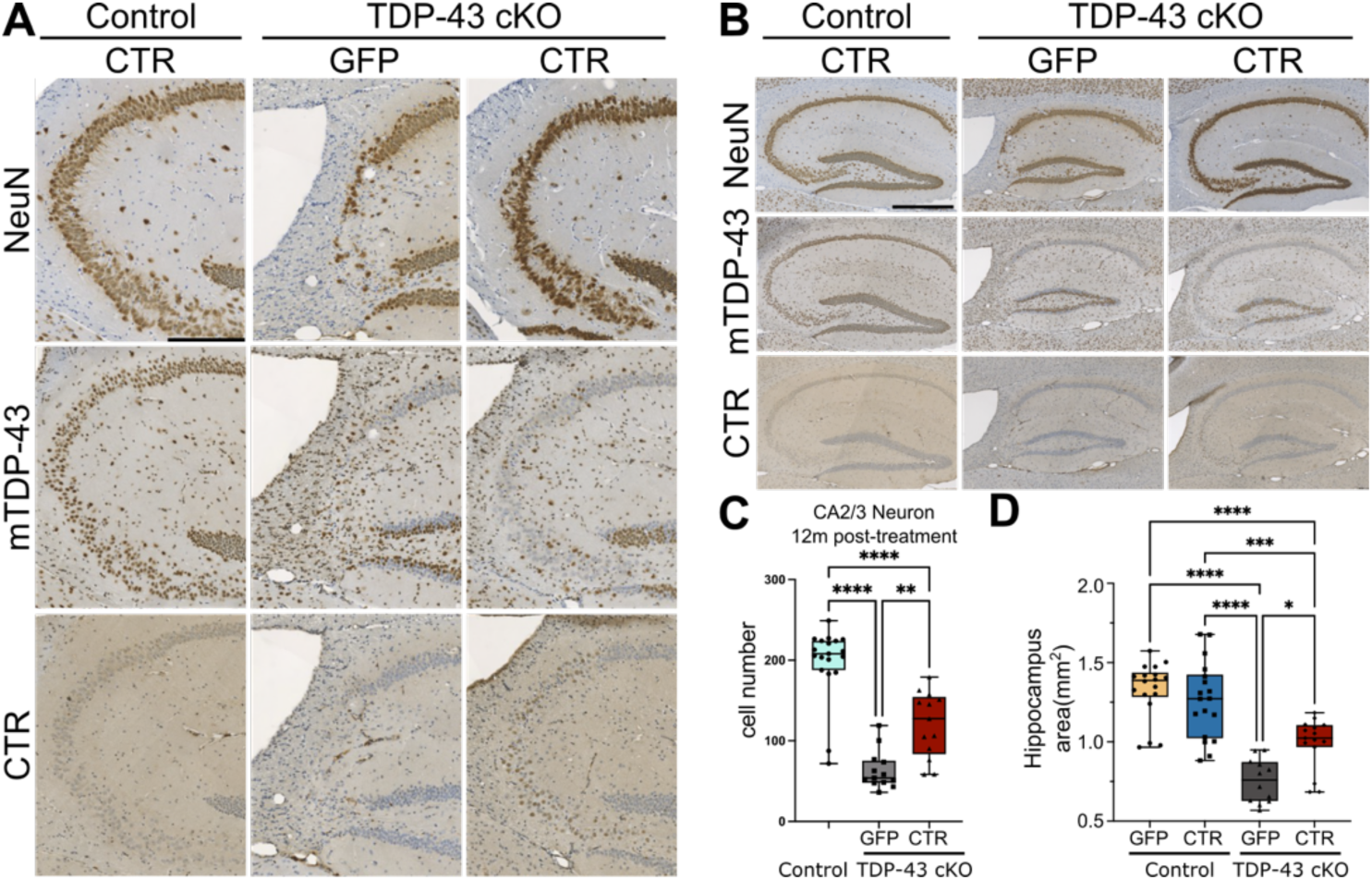
AAV-PHP.eB-CTR mitigates CA2/3 neuron loss and hippocampal atrophy. (A), Representative NeuN immunohistochemistry images of the CA2/3 hippocampal subregion in control and TDP-43 CKO mice 12 months after AAV-CTR or AAV-GFP injection. TDP-43 cKO-GFP mice showed severe neuronal loss in this region, whereas TDP-43 cKO-CTR mice retained more NeuN-positive cells. (B), NeuN immunostaining overview of the entire hippocampus showing global anatomical differences across groups. CTR-treated TDP-43 CKO mice exhibited visibly preserved hippocampal structure. (C), Quantification of NeuN+ neurons in CA2/3 at 12 months post-injection. CTR treatment significantly increased neuron counts in TDP-43 CKO mice compared to GFP controls, though not to the level of control animals. Neuron number in TDP-43 CKO-CTR mice approximated the regional CTR infectivity rate (∼30%)(n = 19 (Control), 13 (TDP-43 cKO GFP), and 13 (TDP-43 cKO CTR) mice. One-way ANOVA, F(2,42) = 48.26, P < 0.0001; Tukey’s post hoc test: Control vs. TDP-43 cKO GFP, P < 0.0001; Control vs. TDP-43 cKO CTR, P < 0.0001; TDP-43 cKO GFP vs. TDP-43 cKO CTR, P = 0.0014.). (D), Quantification of total hippocampal area based on NeuN-stained sections. Control GFP (n = 18) and Control CTR (n = 18) groups had significantly larger hippocampal areas than TDP-43 cKO GFP mice (n = 12) (P < 0.0001). CTR treatment (n = 15) significantly increased hippocampal area in cKO mice compared with GFP-treated cKO animals (P < 0.0001), approaching control group values. All images correspond to 12-month timepoints. Scale bars: a: 200 μm, c: 500 μm. Error bars represent mean ± s.e.m. P-values were calculated using one-way ANOVA followed by Tukey’s multiple comparisons test unless otherwise noted. n values and statistical details are provided in Methods.

Importantly, long-term AAV-PHP.eB-CTR exposure to *Tardbp^f/f^* mice—where TDP-43 expression remains normal—did not lead to any overt phenotypes, including reduction in hippocampal neuron number or area as compared to those observed in untreated or AAV-PHP.eB-GFP treated *Tardbp^f/f^* control littermates. These findings support the idea that AAV-PHP.eB-CTR is highly efficacious, well-tolerated and non-toxic, highlighting its safety profile as a promising therapy to be evaluated for its clinical impact in FTLD-TDP patients.

To evaluate the temporal dynamics and durability of CTR-mediated neuroprotection, we also quantified CA2/3 neurons at different timepoints—6 months and 12 months post-treatment (**Supplementary figure 3A-C**). At 6 months, CA1 neuron numbers did not differ among groups, whereas CA2/3 neurons were significantly reduced in *CamKIIa-CreER;Tardbp^f/f^* mice relative to Control. This CA2/3 loss at 6 months was blunted by CTR (*CamKIIa-CreER;Tardbp^f/f^* + CTR > *CamKIIa-CreER;Tardbp^f/f^* + GFP; partial rescue toward Control, P=0.0536) At 12 months, CA2/3 neuron numbers in *CamKII-CreER;Tardbp^f/^* + CTR mice were significantly higher compare to in controls (**Fig. 2C**), indicating a clear rescue at this later time point. These findings suggest that neurodegeneration progresses gradually in this model and that AAV-PHP.eB-CTR provides sustained protection across the progression of disease. Notably, the preservation of neurons in AAV-PHP.eB-CTR treated *CamKIIa-CreER;Tardbp^f/f^*mice persisted up to 12 months post-treatment which corresponds to the late adult phase of the mouse, underscoring the long-term efficacy of this therapeutic strategy.

Outside the CA2/3 region, neuronal loss was not consistently observed in other hippocampal subfields in our model (**Supplementary figure 3A, C**). However, we noted a marked reduction in overall hippocampal size in *CamKIIa-CreER;Tardbpf/f* mice treated with AAV-PHP.eB-GFP as compared to*Tardbp^f/f^* control littermates. Morphometric analysis of CV-stained sections confirmed a significant rescue of hippocampal area in the AAV-PHP.eB-CTR treated group (**Fig. 2D; Supplementary figure 4**), indicating that CTR expression alleviates TDP-43–dependent neurodegeneration at both cellular and anatomical levels.

### AAV-PHP.eB-CTR rescues social and cognitive deficits

Since we showed behavioral deficits in *CamKIIa-CreER;Tardbpf/f* mice^29^, we asked whether AAV-PHP.eB-CTR treatment rescues behavioral impairments associated with TDP-43 depletion. As before, we conducted a battery of behavioral tests approximately 4-5 months after AAV-PHP.eB-CTR treatment. During the first round of behavioral testing, both male and female *CamKIIa-CreER;Tardbp^f/f^* mice and *Tardbp^f/f^*littermate controls underwent assessments including the open field test, light–dark box test, novel object recognition (NOR) test, and the social behavior test. No significant differences were observed in locomotor activity or anxiety-related behavior between genotypes or treatment groups in the open field and light–dark box tests(**Supplementary figure 5A-E**). However, deficits emerged in the novel object and social novelty preference paradigms. In the NOR test, *CamKIIa-CreER;Tardbp^f/f^*mice with AAV-PHP.eB-CTR or control AAV-PHP.eB-RFP treatment spent equal time exploring familiar and novel objects, indicating impaired object recognition memory. In contrast, *CamKIIa-CreER;Tardbp^f/f^* mice treated with AAV-PHP.eB-CTR displayed restored object novelty preference, like *Tardbp^f/f^* control littermates (**Fig. 3A, D**).

**Figure 3.**
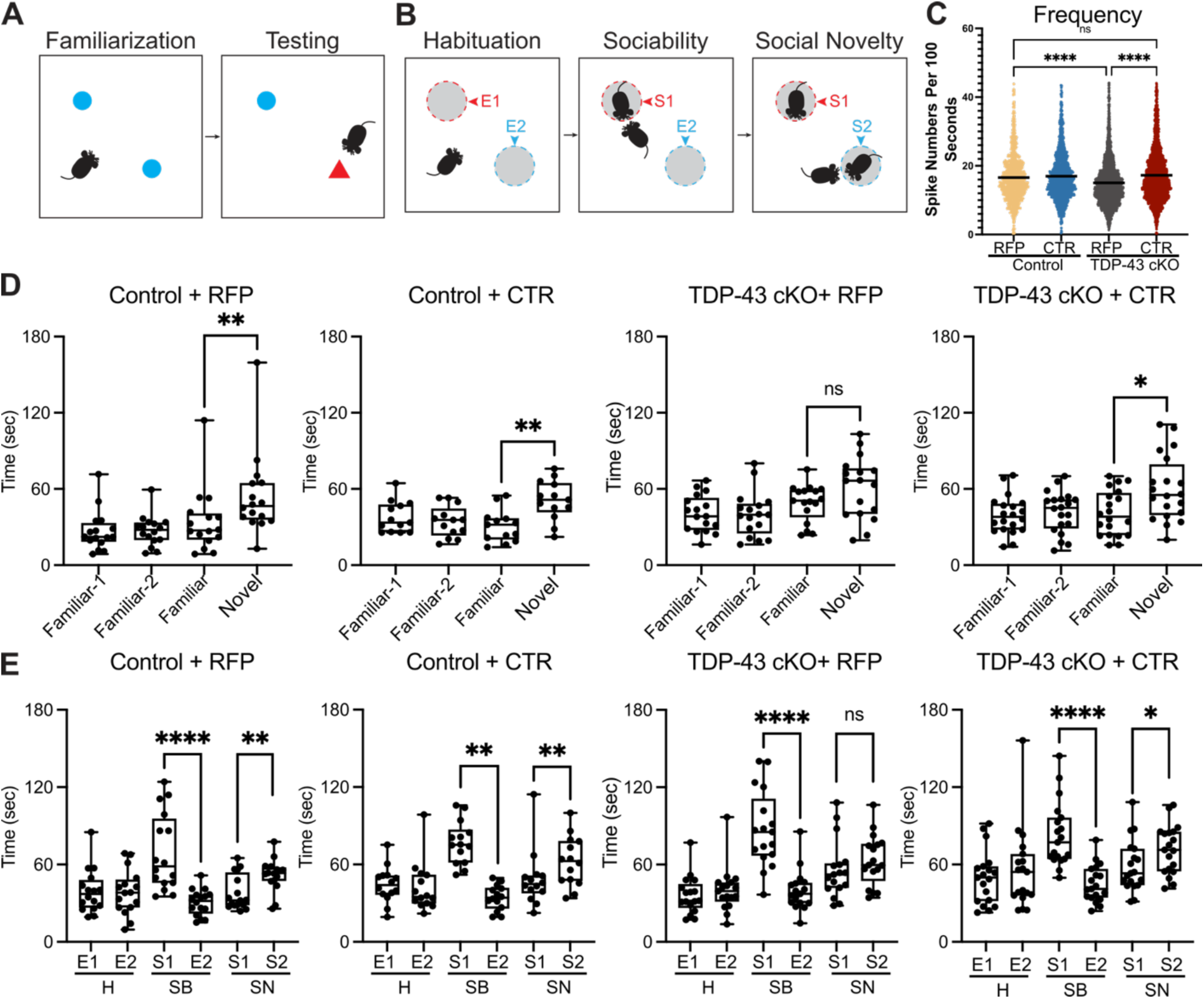
AAV-PHP.eB-CTR restores neuronal calcium activity and attenuates memory deficits. (A), Schematic of the novel object recognition (NOR) test protocol. Mice were exposed to two identical objects during a familiarization phase, followed by a testing phase in which one object was replaced with a novel item. Time investigating each object was measured. (B), Schematic of the social behavior test. Mice underwent three 10-minute phases: habituation (two empty cages), sociability (one unfamiliar mouse introduced), and social novelty (second unfamiliar mouse added). (C), Analysis of *in vivo* neuronal activity using calcium imaging in the prelimbic cortex. Compared to Control-RFP controls, TDP-43 cKO-RFP mice exhibited reduced neuronal calcium spike frequency, consistent with impaired excitatory activity. CTR treatment normalized activity levels. Imaging was performed through GRIN lenses following AAV-GCaMP6f injection. Group sizes: Control-RFP (n = 936 ROIs), Control-CTR (n = 1379 ROIs), TDP-43 cKO-RFP (n = 2285 ROIs), TDP-43 cKO-CTR (n = 2334 ROIs). **** indicates P < 0.0001, ns = not significant. (D), Quantification of exploration time during the novel object recognition (NOR) test. TDP-43 cKO-RFP mice (n = 17) failed to show a preference for the novel object, indicating impaired recognition memory. CTR treatment (n = 20) restored novel object preference to levels comparable to Control-RFP (n = 16) and Control-CTR (n = 13) groups. (E), Quantification of time spent investigating social targets. While all groups displayed normal sociability, TDP-43 cKO-RFP mice failed to show a preference for social novelty. CTR-treated TDP-43 cKO mice showed restored preference for the novel conspecific, indicating rescue of social memory. Data are mean ± s.e.m. P-values were calculated using Kruskal–Wallis ANOVA followed by Dunn’s post hoc tests or Mann–Whitney tests, as noted in Methods. ****P<0.0001, **P<0.01, *P<0.05, ns = not significant.

A comparable rescue pattern was observed in the social behavior test. During the sociability phase, all groups exhibited normal preference for a novel conspecific over an empty enclosure. However, during the social novelty phase, *CamKIIa-CreER;Tardbp^f/f^*mice treated with AAV-PHP.eB-RFP failed to show a preference for a second novel mouse over the relatively familiar conspecific, consistent with impaired social recognition. This behavioral deficit was rescued in AAV-PHP.eB-CTR treated *CamKIIa-CreER;Tardbp^f/f^* mice, who displayed normal social novelty preference, comparable to *Tardbp^f/f^* control animals (**Fig. 3B, E**). These results indicate that AAV-PHP.eB-CTR treatment reverses specific cognitive and social deficits associated with TDP-43 loss in forebrain neurons.

### AAV-PHP.eB-CTR restores neuronal calcium activity impaired by TDP-43 loss

Following the initial behavioral assessments, mice underwent brain surgeries including viral injection of AAV-CaMKII-GCaMP6f and GRIN lens implantation targeting the prelimbic cortex to enable longitudinal *in vivo* calcium imaging. Perioperative outcomes are summarized in Supplementary Table 1: 7 of 41 male mice (17%) and 5 of 26 female mice (19%) died during or shortly after surgery. Of the 67 mice subjected to both procedures, 30 (45%) yielded high-quality calcium imaging data, with each experimental group represented.

To quantify neuronal activity, 15-minute calcium imaging recordings were processed using a custom Python and MATLAB pipeline. The firing frequency of active neurons was extracted and pooled across sexes for analysis. TDP-43 LOF in *CamKIIa-CreER;Tardbp^f/f^* +RFP mice significantly reduced the frequency of calcium transients compared to that of *Tardbp^f/f^* control littermates (**Fig. 3C**), indicating functional impairment of excitatory neuronal activity. Remarkably, AAV-PHP.eB-CTR treatment restored calcium firing frequency to levels comparable to those observed in *Tardbp^f/f^* control mice (**Fig. 3C**). These results suggest that AAV-PHP.eB-CTR rescues not only behavioral deficits, but also forebrain circuit abnormalities.

Together, these results demonstrate that AAV-PHP.eB-CTR not only corrects molecular splicing defects but also preserves forebrain neuronal circuit activity and restores behavioral performance, supporting its therapeutic potential for testing in the clinic for FTLD-TDP.

## Discussion

Our findings establish the success of a chimeric splicing repressor (CTR) designed to restore TDP-43’s splicing repression activity *in vivo* while avoiding its aggregation. Using a conditional TDP-43 knockout mouse model that mimics the early stage of FTLD-TDP, we demonstrate that ICV delivery of AAV-PHP.eB-CTR resulted in broad repression of cryptic exon inclusion, long-term rescue of neuronal survival in vulnerable hippocampal neurons, and restoration of cognitive behavior and neuronal calcium dynamics. These findings validate CTR as a disease-modifying payload capable of restoring key functional aspects of TDP-43 in the adult forebrain and provide preclinical proof-of-concept for an AAV serotype amenable for broad biodistribution to the adult brain designed to restore TDP-43 function for C9orf72-FTD or FTLD-TDP.

Current therapeutic strategies aimed at mitigating TDP-43 loss-of-function primarily rely on antisense oligonucleotides (ASOs) designed to suppress individual cryptic exons, such as those found in *STMN2* and *UNC13A*^19,40–42^. While these approaches have demonstrated transcript-level correction and partial phenotypic rescue in preclinical models, they are inherently constrained in scope. Each ASO targets a single downstream consequence of TDP-43 dysfunction, necessitating multiple independent interventions to address complex disease phenotypes^43,44^. This piecemeal approach may be insufficient to restore network-level RNA homeostasis in the affected neurons.

In contrast, CTR represents a unified strategy that intervenes at the level of the splicing repression mechanism itself. Rather than correcting the results of TDP-43 dysfunction one transcript at a time, CTR restores the upstream regulatory function that governs a broad range of cryptic exon inclusion, as well as other functions such as alternative polyadenylation. This distinction not only provides therapeutic breadth but also aligns more closely with the pathophysiology of the disease.

Moreover, whereas full-length TDP-43 re-expression poses risks due to its aggregation-prone domains and overexpression toxicity, CTR is structurally designed to mitigate these issues^1,44^. CTR avoids reintroduction of the aggregation-prone C-terminal domain of TDP-43, which has been implicated in pathological phase separation and neurotoxicity^45^. Moreover, by retaining the 3′UTR of human *TARDBP*, CTR preserves autoregulatory feedback, minimizing the risk of overexpression—a critical safety feature especially in the context of viral gene delivery^39^. By combining mechanistic precision with reduced off-target liability, CTR offers a complementary and potentially more versatile alternative to ASO-based interventions—particularly in disorders like FTLD-TDP or C9-FTD where multiple gene networks are disrupted in parallel.

Finally, this demonstrated efficiency of AAV-PHP.eB in CNS-wide transduction of adult mice highlights the potential of developing new BBB permeable AAV serotypes to enable efficient delivery of payload to the human adult brain^46,47^ to include other neurodegenerative disorders exhibiting pathology of TDP-43 such as LATE^48^, and AD-TDP^49–51^ Moreover, the therapeutic efficacy of CTR appears tightly linked to local rate of transduction, emphasizing the need for improved delivery platforms for clinical application—such as next-generation capsids with broader tropism and non-invasive administration routes^52,53^.

## Methods

### Animal Models and Genotyping

All animal procedures were approved by the Institutional Animal Care and Use Committee (IACUC) at Johns Hopkins University or University of Wyoming and conducted in accordance with NIH guidelines. To model TDP-43 loss of function in a cell-type– specific manner, we utilized a previously established conditional Tardbp knockout mouse line (Tardbp^f/f^), in which exon 3 of the Tardbp gene is flanked by LoxP sites^54^(The Jackson Laboratory, stock #017591).We crossed these mice with the *CamkIIa*-CreERT2 driver line to generate a tamoxifen-inducible, excitatory neuron–specific Tardbp knockout model^55,56^ (hereafter referred to as TDP-43 cKO).

Experimental cohorts included Tardbp^f/f^; *CamkIIa*-CreERT2 mice (referred to as TDP-43 CKO) and Tardbp^f/f^ littermates lacking Cre (referred to as Control). Both sexes were used, and groups were balanced by age and sex. Recombination was induced by feeding mice tamoxifen-containing chow (Envigo, TD.130859, 400 mg/kg) for 4 weeks, followed by a 2-week washout period.

Mice were housed in a 12-hour light/dark cycle with ad libitum access to food and water, and group-housed when possible. Genotyping was performed using DNA extracted from ear biopsies. PCR was conducted using primers specific to the loxP-flanked Tardbp allele and the Camk2a-CreERT2 transgene: PCR products were analyzed by agarose gel electrophoresis. Genotyping Primers are as follows:

*CamKIIa*-Cre F: GACAGGCAGGCCTTCTCTGAA

*CamKIIa*-Cre R: CTTCTCCACACCAGCTGTGGA

*Tardbp* FF F: AACTTCAAGATCTGACACCCTCCCC

*Tardbp* FF R: GGCCCTGGCTCATCAAGAACTG;

The expected product of the *CamkIIa*-Cre primers is 536bp. The expected products of the *Tardbp* FF primers are 376 bp for the floxxed product and 230 bp for the wild-type product.

### AAV Vector Design and Delivery

The therapeutic construct CTR (Chimeric TDP-43 Repressor) was designed by fusing the N-terminal RNA recognition domains (RRM1 and RRM2, residues 1–267) of human TDP-43 with the splicing repressor domain of RAVER1 (residues 450–643), replacing the aggregation-prone C-terminal glycine-rich region^1^. The construct includes the endogenous *TARDBP* 3′ untranslated region (3′UTR) to preserve TDP-43 autoregulatory feedback. The CTR transgene was driven by the CBh (chicken β-actin hybrid) promoter.

Control vectors included AAV9-CMV-GFP (Cat# AAV-0090, Virovek), with a reported titer of 2E13 vg/mL, and AAV9-CMV-mCherry, both obtained from the same supplier.

For adult intracerebroventricular (ICV) delivery, mice were anesthetized with isoflurane and placed in a stereotaxic frame. A total of 5 μL (1E11 vg) was injected into the right lateral ventricles at the following coordinates from bregma: AP –0.5 mm, ML 1.0 mm, DV –2.5 mm^57^. Injection was performed at a rate of 1 μL/min using a 5 μL Hamilton syringe. Mice were allowed to recover on a heating pad and monitored until fully ambulatory.

For neonatal injections, pups were injected within 8 h of birth. Cryo-anesthesia was induced by brief exposure to wet ice (<2 min) with a protective barrier to prevent skin injury. A pulled glass capillary needle (41 mm, ∼0.5 mm tip) was inserted ∼3 mm into the lateral ventricle, and 3 μL of AAV9 (1×10^13 vg/mL) encoding either CTR or GFP was slowly delivered over ∼15 s. Trypan blue (0.05%) was included in the injection solution to aid visualization. Following injection, pups were rewarmed under a heat source, placed in bedding from the home cage to restore maternal scent, and returned to the dam once active. Survival was ∼80%, with early losses primarily due to incomplete recovery from anesthesia (P0–P1) and occasional hydrocephalus developing at later stages (P14–P21). Pups showing signs of distress were euthanized according to IACUC guidelines.

### Tissue Collection and Immunohistochemistry

Mice were euthanized at designated timepoints (1-, 3-, 6-, or 12-months post-injection) by isoflurane overdose, followed by transcardial perfusion with phosphate-buffered saline (PBS) and then 4% paraformaldehyde (PFA) in PBS. Brains were carefully dissected and post-fixed in 4% PFA at 4°C overnight, then dehydrated, embedded in paraffin, and sectioned sagittally at 10 μm thickness using a rotary microtome (Leica HistoCore MULTICUT).

For immunohistochemistry, slides were first oven-baked at 60°C for 30 mins, then deparaffinized in xylene and rehydrated through a graded ethanol series (100%, 95%) to water. Antigen retrieval was performed in 10 mM sodium citrate buffer (pH 6.0) by heating slides in a microwave for 4 minutes. After cooling, sections were encircled with a hydrophobic barrier pen and incubated in blocking solution containing 1.5% normal goat serum and 0.1% Triton X-100 in PBS for 1 hour at room temperature. Primary antibodies diluted in blocking buffer were applied overnight at 4°C in a humidified chamber. CTR expression in mouse was expressed using N-terminal targeting human TDP-43 antibody. Primary antibodies used:

human-TDP-43 (Ref#WH0023435M1, 1:1000, Millipore Sigma),

mouse-specific TDP-43 (C-Terminus, Ref#12892-1-AP, 1:1000, Proteintech),

Mouse-NeuN (Cat#MAB377,1:2000, MERCK).

Endogenous peroxidase activity was quenched by incubating slides in 0.3% hydrogen peroxide (H₂O₂) in methanol for 30 minutes, followed by three 10-minute washes in PBST (PBS + 0.1% Tween-20). Slides were incubated with biotinylated secondary antibodies (Vector Laboratories, BP-9100-50, BP-9200-50) for 1 hour at room temperature. Signal was amplified using VECTASTAIN Elite ABC reagent (Vector Laboratories, PK-7100) for 1 hour and visualized by DAB substrate reaction (Vector DAB Peroxidase Substrate Kit, SK4100), monitoring under a microscope for optimal development. Slides were then counterstained with hematoxylin, dehydrated through graded alcohols, cleared in xylene, and mounted with mounting media.

Brightfield images were captured using a ZEISS microscope equipped with a high-resolution digital camera. Quantification was performed using ImageJ with uniform region-of-interest (ROI) definitions applied across experimental groups.

### Reverse Transcription PCR (RT-PCR)

Total RNA was extracted from dissected hippocampal tissue using the RNeasy Mini Kit (Qiagen, #74106) according to the manufacturer’s protocol. RNA concentration and purity were assessed using a Nanodrop spectrophotometer (Thermo Fisher, 13-400-519), and RNA integrity was confirmed by agarose gel electrophoresis.

First-strand cDNA was synthesized using the ProtoScript® First Strand cDNA Synthesis Kit (New England Biolabs, E6300) following the manufacturer’s protocol. Each reaction used 500 ng to 1 μg of total RNA and was primed with a mixture of enzyme mix and random primers to ensure coverage of all transcripts.

To assess cryptic exon inclusion, we designed primers flanking known TDP-43– regulated cryptic exons (e.g., *Adnp2, Ap3b2, Bud23, Camk1g, Crem, Ggct, Unc13a, Synj2bp, Tbc1d1, Usp15, Tecpr1,* and *Washc4*)^30^. Products were separated on 1.5% agarose gels and visualized using GelRed staining under UV illumination. Band intensities were quantified using ImageJ and normalized to total transcript signal (i.e. inclusion + exclusion bands).

To evaluate CTR autoregulation, a primer pair was designed targeting the N-terminal human TDP-43 sequence (RRM1) and the C-terminal RAVER1 fusion domain. Amplification of this junction-specific product confirmed the presence of CTR transcript.

All PCR reactions were performed in technical duplicates or triplicates, and at least three biological replicates per condition were included. Touch down thermal cycling conditions were optimized per primer set and are provided in Supplementary Table 2-4.

### BaseScope In Situ Hybridization

BaseScope™ in situ hybridization was performed using the BaseScope™ v2 Assay (Advanced Cell Diagnostics, ACD) according to the manufacturer’s protocol. Paraffin-embedded sagittal brain sections (10 μm thickness) were processed using standard deparaffinization, target retrieval, and protease digestion steps as described in the BaseScope manual.

Probe sets were custom-designed by ACD to target exon-exon junctions spanning cryptic exon inclusion events. Specifically, we used a probe targeting the *Unc13a*(CAT#1182491-C1), *Synj2bp*(CAT#712191), and *Ift81*(CAT#712201) cryptic exons along with one targeting CTR mRNA(CAT#1573691-C1). Signal amplification and chromogenic detection were carried out using the BaseScope™ Red Detection Kit. Slides were counterstained with 50% hematoxylin, air-dried, and mounted using AquaMount.

Brightfield images were acquired using a ZEISS microscope. For quantification, the number of BaseScope puncta per nucleus was manually counted within defined hippocampal regions of interest using ImageJ. At least three sections per mouse and 3– 5 mice per group were analyzed in a blinded fashion.

### Behavioral Testing

Behavioral experiments were conducted at the University of Wyoming Animal Behavior Core Facility under blinded conditions. All testing was performed during the light phase (9:00 a.m. to 5:00 p.m.), and mice were habituated to the testing room for at least 30 minutes prior to each assay.

Open field testing was used to evaluate general locomotion and anxiety-like behavior. Mice were placed in a 42 × 42 cm open-field arena for 15 minutes. Distance traveled and time spent in the center zone were tracked using Noldus EthovisionXT 15 software.

Novel object recognition (NOR) was used to assess recognition memory. Mice were exposed to a 42 × 42 cm open-field arena settled with two identical objects for 10 minutes. After a 5-minutes delay, the arena was cleaned and one object was replaced with a novel object of similar size, and mice were reintroduced to the arena for 10 minutes. Object exploration was scored manually by trained observers blinded to group allocation.

Social behavior testing was performed and recorded in an adapted same-chambered procedure as previously described^58^. During habituation, mice were allowed to freely explore a 42 x 42 cm arena settled with two empty wire cages at the opposite corners for 10 minutes. In the 10-minutes sociability test phase, an unfamiliar age- and sex-matched conspecific (stranger 1) was enclosed in one of the empty wire cages. In the 10-minutes social novelty test phase, a second unfamiliar age- and sex-matched conspecific (stranger 2) was placed in the previously empty wire cage. Time spent sniffing or touching each cage were scored manually frame by frame by trained observers blinded to group allocation.

All behavioral assessments were performed by experimenters blinded to genotype and treatment group. Data were analyzed as described in the statistical analysis section.

### *In vivo* calcium imaging

*In vivo* calcium imaging was performed to monitor neuronal activity in the prelimbic cortex of TDP-43 cKO and control mice. 500nl of AAV1-CamKII-GCaMP6f (Addgene, 2 E13 GC/mL) was injected stereotaxically into the prelimbic cortex (coordinates: A/P +1.9 mm, M/L 0.5 mm, D/V 1.75 mm) under isoflurane anesthesia as previously described^59^. A 1 mm diameter GRIN lens (Grintech) was implanted into the injection site to reach the depth of 1.8 mm and secured with dental cement as previously described^59^. Mice were allowed to recover for at least 4 weeks prior to recording.

*In vivo* Ca^2+^ Imaging was performed using a previously established custom-built miniature fluorescence microscope recording system^60^ during exploratory behavior in a 42 x 42 cm open-field arena. Three recording sessions, each lasting 5 minutes per mouse were conducted. Calcium fluorescence signals were acquired at 10 Hz and preprocessed using standard motion correction and ΔF/F0 normalization. Individual neurons were identified and segmented using CNMF-E implemented in MATLAB. Calcium event rates were quantified by thresholding deconvolved signals, and the average spike rate per neuron was calculated for each animal as previously described^61^.

Mice were excluded from analysis if recordings lacked sufficient signal-to-noise or spatially stable fields of view. Of 67 mice that underwent imaging procedures, 30 (45%) produced data of sufficient quality for analysis (see Supplementary Table 1 for group breakdown). Sample sizes per group ranged from 5 to 9 animals. All animals were included in downstream group-level comparisons.

## Data analysis

All statistical analyses were performed using GraphPad Prism (version 10) unless otherwise specified. Data are presented as mean ± standard error of the mean (s.e.m.), unless otherwise indicated. Group comparisons were evaluated using unpaired two-tailed Student’s t-tests, Mann-Whitney U test, one-way ANOVA, or two-way ANOVA with Holm– Sidak post hoc correction, depending on experimental design and variance structure.

## Acknowledgements

This work was supported in part by the National Institutes of Health grants R01 NS095969 (to P.C.W.), UG3/UH3 NS115608 (to P.C.W.), R33NS115161 (to P.C.W.), R01 NS129878 (to P.C.W. and Y.L.), and the Intramural Research Program of the National Institutes of Health (NIH). The contributions of the NIH author (D.-T.L.) are considered Works of the United States Government. The findings and conclusions presented in this paper are those of the author(s) and do not necessarily reflect the views of the NIH or the U.S. Department of Health and Human Services.

## Author contributions

T.C. and P.C.W. conceptualized, designed and interpreted the study. T.C., Y.L, J.P.L. and P.C.W. wrote the manuscript. T.C., R.T., R.L., A.P.M., I.R.S, M.S.B, G.D.B., B.P. and X.W. performed experiments. D.-T.L. provided the custom-build miniscope recording system. All the authors reviewed and approved the final manuscript.

## Supplementary data figure legends

**Supplementary figure 1.**
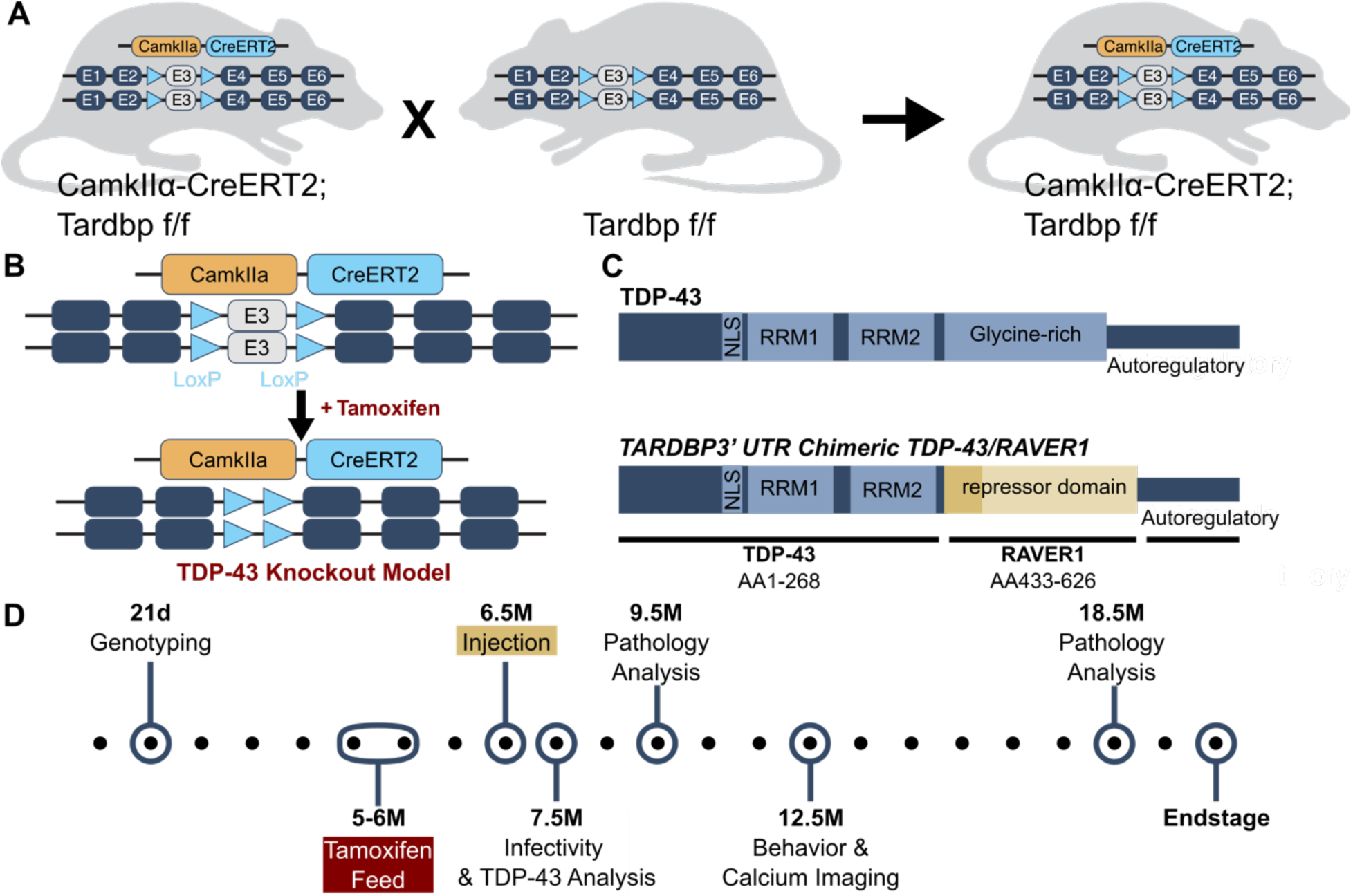
Conditional TDP-43 knockout model and AAV-CTR vector therapeutic design. (**A,B**), Schematic of the inducible, excitatory neuron–specific *Tardbp* conditional knockout (TDP-43 CKO) mouse model. LoxP sites flank exon 3 of *Tardbp* in *Tardbp*^f/f^ mice, and CreERT2 is driven by the Camk2a promoter. Upon tamoxifen administration, exon 3 is excised in forebrain excitatory neurons, resulting in loss of functional TDP-43. (C), Diagram of the CTR construct (Chimeric TDP-43 Repressor), composed of the N-terminal RNA recognition motifs (RRM1 and RRM2) of TDP-43 (amino acids 1–267), fused to the splicing repression domain (RAVER1, amino acids 450–643), and followed by the endogenous human *TARDBP* 3′ untranslated region (3′UTR) to preserve autoregulatory feedback. (D), Timeline of the *in vivo* study design. Mice were fed tamoxifen at 5 months of age for 1 month to induce recombination. After a 2-week recovery, intracerebroventricular injection of AAV-PHP.eB vectors expressing CTR or GFP was performed. Mice were analyzed at 1-, 3-, 6-, and 12-months post-injection.

**Supplementary figure 2.**
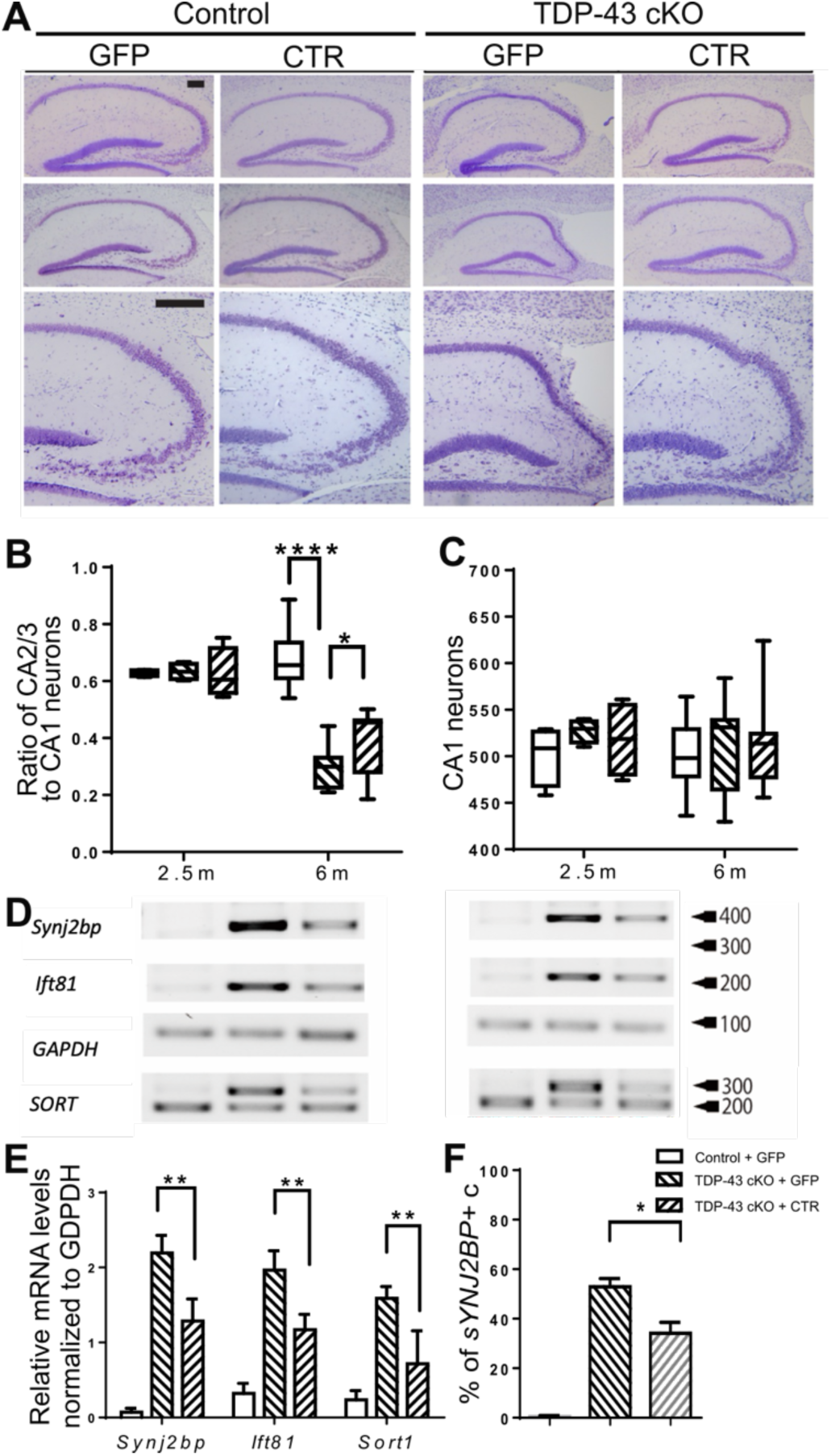
AAV9-ICV Delivery of CTR at P0 Attenuates Cryptic Exon Inclusion of TDP-43 Target Genes and Preserves Neuronal Integrity in the Hippocampus. **(A)**, Representative cresyl violet staining of the hippocampus in Control + GFP, Control + CTR, TDP-43 cKO + GFP, and TDP-43 cKO + CTR mice at 2.5 and 6 months of age. **(B),** Quantification of the ratio of neurons in the CA2/3 region, normalized to neuron numbers in CA1 region, 2.5 m time point: Control + GFP (n = 4), TDP-43 cKO + GFP (n = 4), and TDP-43 cKO + CTR (n = 4), 6 m time point: Control + GFP (n = 9), TDP-43 cKO + GFP (n = 9), and TDP-43 cKO + CTR (n = 12) **(C)**, Quantification of neurons in the CA1 region in the same groups as panel B. **(D),** Representative semi-quantitative RT-PCR analysis of three TDP-43 target cryptic exon RNAs (Synj2bp, Ift81, and Sort) in hippocampus and cortex of Control + GFP (n = 3), TDP-43 cKO + GFP (n = 4), and TDP-43 cKO + CTR (n = 3) mice at 4 months of age. **(E)**, Quantification of cryptic Synj2bp, Ift81, and Sort RT-PCR cryptic products in hippocampus from the same groups as in (d). **(F)**, Quantification of cryptic Synj2bp positive cells by *In situ* hybridization in hippocampus from the same groups. Band intensities were quantified using ImageJ. One-way ANOVA: **** P < 0.0001, ## P < 0.01. Scale bars: 200um.

**Supplementary figure 3.**
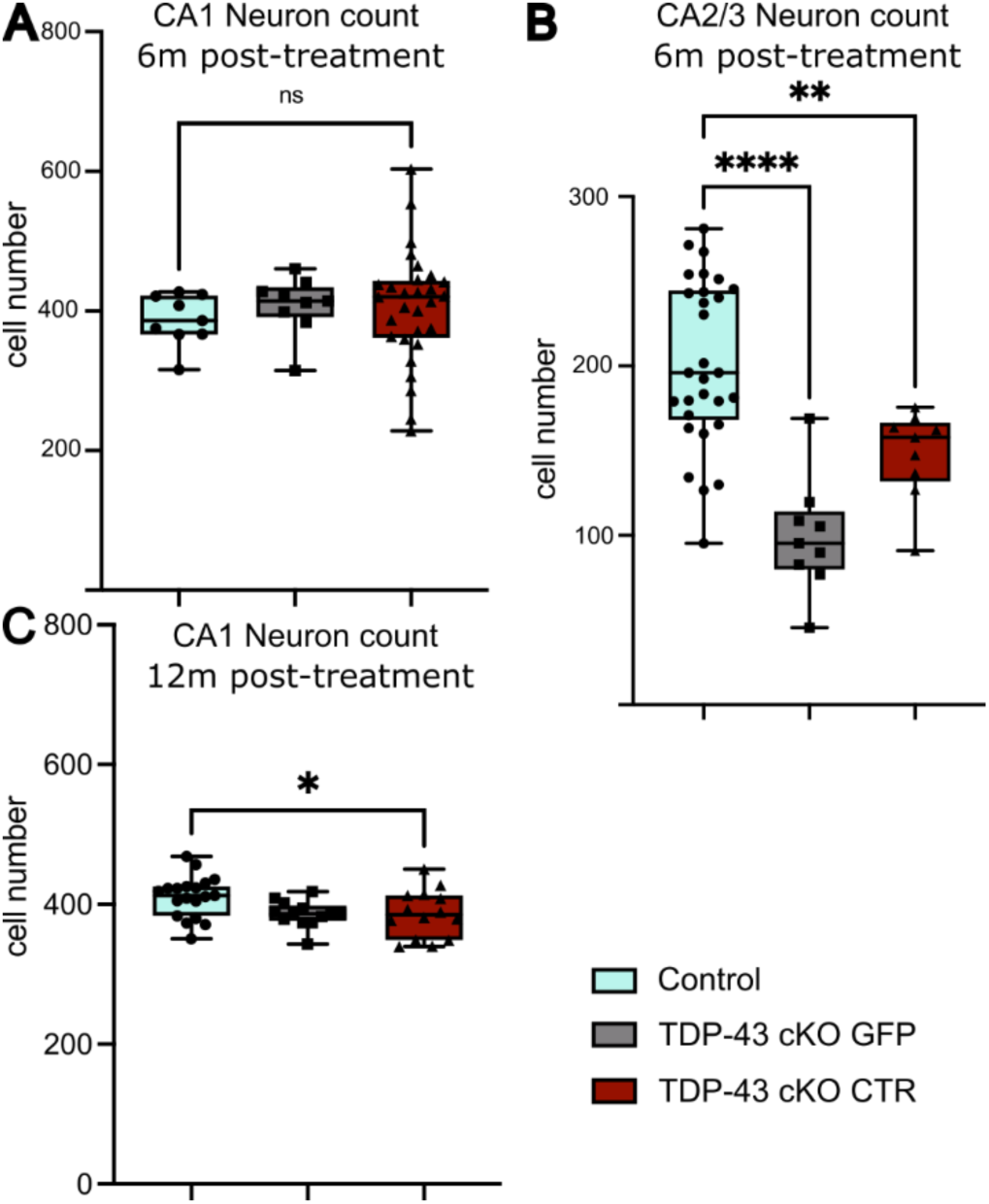
Neuronal survival after CTR Treatment. (A) Quantification of CA1 neuron numbers at 6 months post-treatment in Control(n=29), TDP-43 cKO + GFP(n=9), and TDP-43 cKO + CTR(n=9) mice. No significant differences were observed between groups. (B) Quantification of CA2/3 neuron numbers at 6 months post-treatment in the same groups as in (a). TDP-43 cKO + GFPmice showed a significant reduction compared to Control, and CTR expression partially rescued neuron numbers. (C) Quantification of CA1 neuron numbers at 12 months post-treatment in in Control(n=19), TDP-43 cKO + GFP(n=13), and TDP-43 cKO + CTR(n=13) mice. Data are presented as mean ± SD. Statistical analysis was performed using one-way ANOVA followed by post hoc multiple comparisons. Statistical significance: p < 0.05 (*), p < 0.01 (**), p < 0.0001 (***), ns = not significant.

**Supplementary figure 4.**
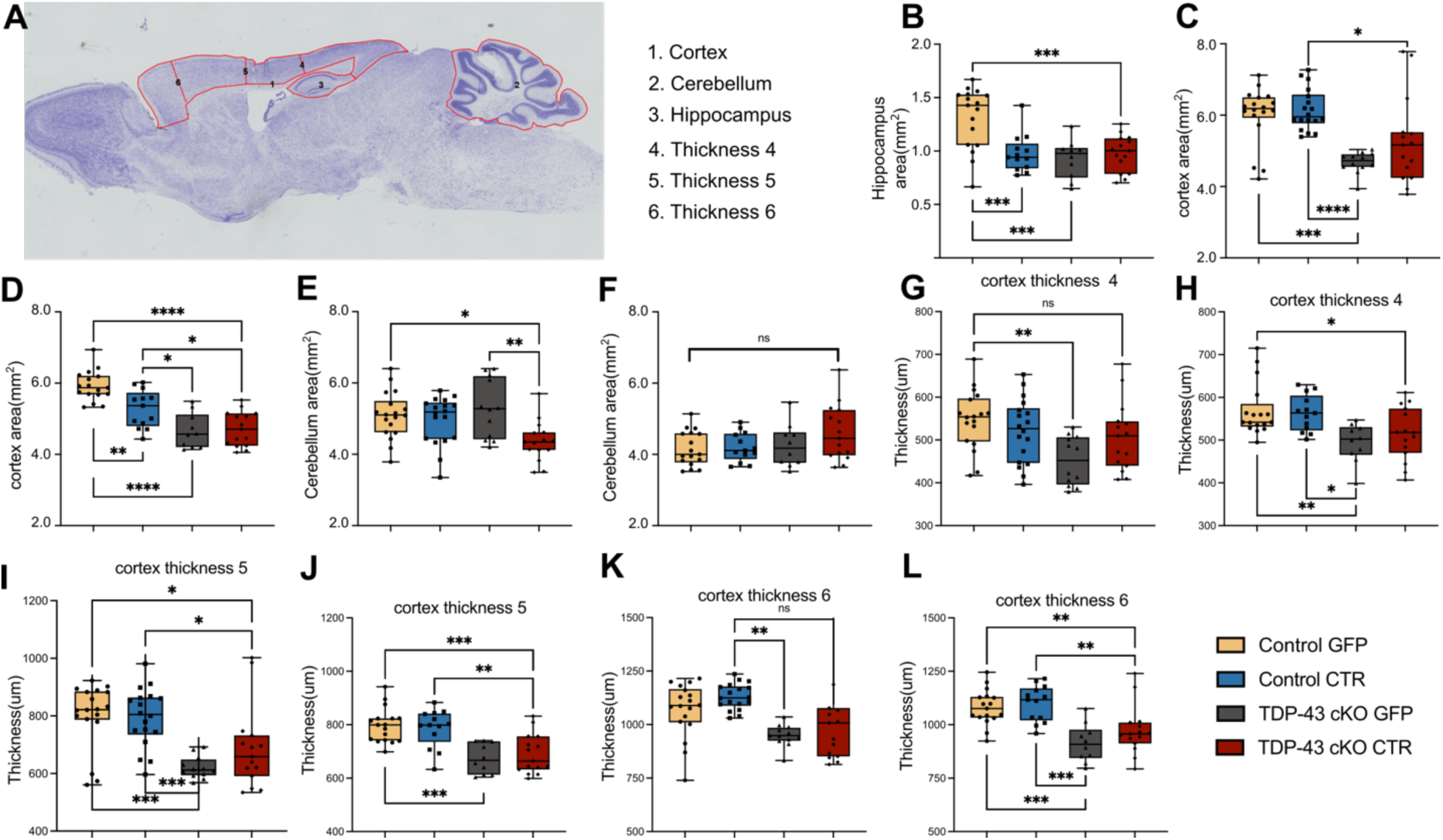
Morphometric analysis of cortical thickness, cerebellar area, and hippocampal area after CTR treatment in Control and TDP-43 cKO mice. (A) Schematic illustration of the brain regions measured for cortical thickness and area, cerebellar area, and hippocampal area. Red outlines indicate the regions of interest (ROIs): cortex (1, 4–6), cerebellum (2), and hippocampus (3). (**G, I, K**) Quantification of cortical thickness in the indicated cortical regions at 6 months after treatment across experimental groups: Control + GFP(n=10), Control + CTR(n=15), TDP-43 cKO + GFP(n=16), and TDP-43 cKO + CTR(n=13). (**H, J, L**) Quantification of cortical thickness in the indicated cortical regions at 12 months after treatment across experimental groups: Control + GFP(n=18), Control + CTR(n=18), TDP-43 cKO + GFP(n=12), and TDP-43 cKO + CTR(n=15). (**B, D, F**) Quantification of cerebellar area (F), cortical area (D), and hippocampal area (B) at 6 months after treatment. (**C, E**) Quantification of cerebellar area (E) and cortical area (C) at 12 months after treatment. Data are presented as mean ± SD. Statistical analysis was performed using one-way ANOVA followed by post hoc multiple comparisons. Statistical significance: p < 0.05 (*), p < 0.01 (**), p < 0.0001 (***), ns = not significant.

**Supplementary figure 5.**
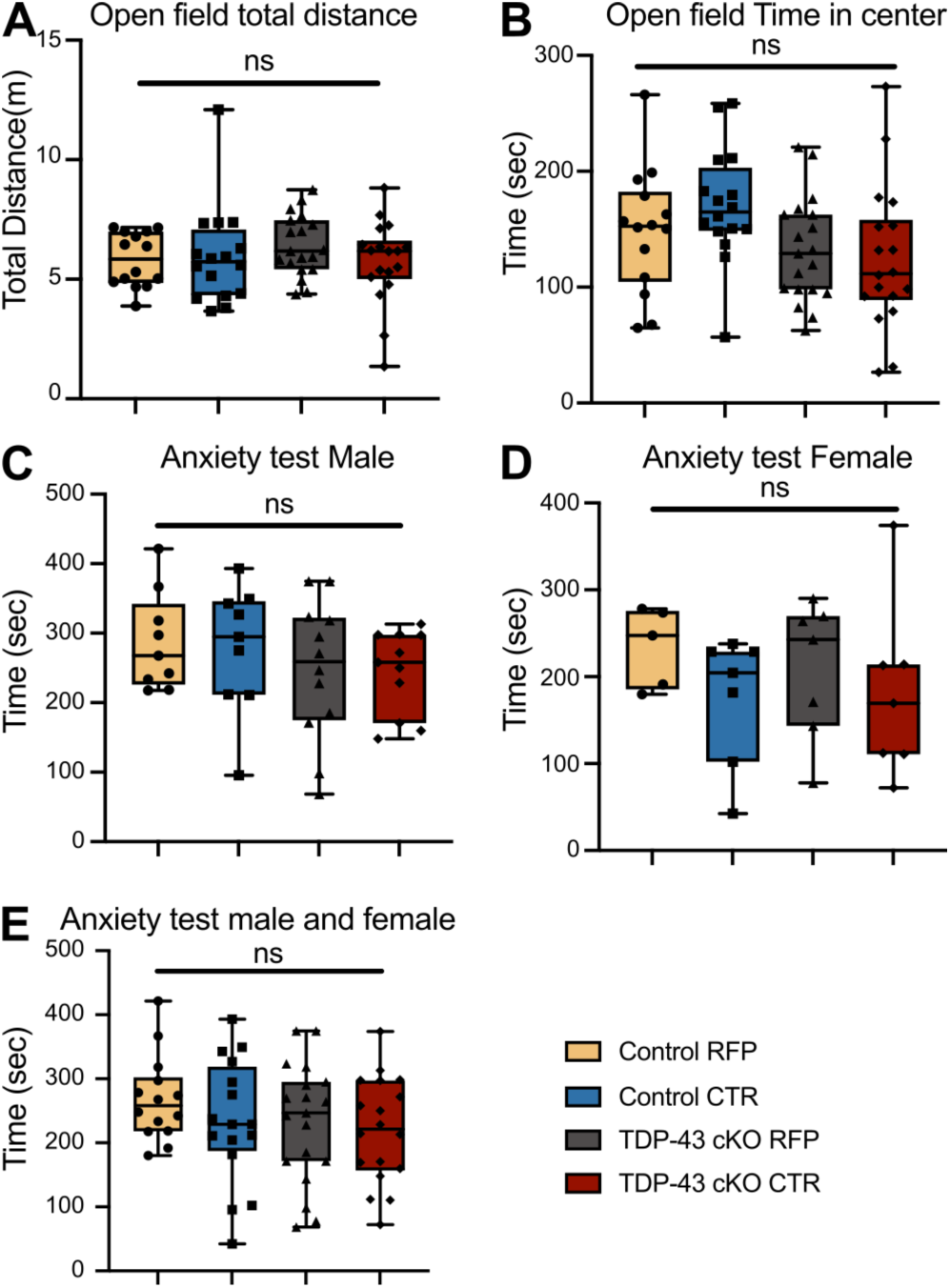
No significant differences were observed in locomotor activity or anxiety-related behavior between genotypes or treatment groups in the open field and light–dark box tests. **(A)** Total distance traveled in the open field test. **(B)** Time spent in the center of the open field arena. **(C)** Time spent in the light compartment of the light–dark box test in male mice. **(D)** Time spent in the center of the open field in female mice. **(E)** Time spent in the light compartment of the light–dark box test in male and female mice combined. Groups: Control + RFP(N = 16), Control + CTR(N = 14), TDP-43 cKO + RFP(N = 18), and TDP-43 cKO + CTR(N = 19). Data are presented as mean ± SEM; ns = not significant by one-way ANOVA.

## Supplementary Tables

**Supplementary Table 1.** Perioperative mortality and calcium imaging data yield by sex and experimental group.

|  | Male(n =<br>41) | Female(n<br>= 26) | Total(n =<br>67) |
| --- | --- | --- | --- |
| Deaths during/after surgery,<br>n (%) | 7 (17%) | 5 (19%) | 12 (18%) |
| Mice with high-quality<br>calcium imaging data, n (%) | 13 (32%) | 17 (65%) | 30 (45%) |
| Breakdown of mice with high-quality data |  |  |  |
| Control + CTR | 3 | 4 | 7 |
| Control + RFP | 3 | 3 | 6 |
| TDP-43 cKO + CTR | 2 | 4 | 6 |
| TDP-43 cKO + RFP | 5 | 6 | 11 |

**Supplementary Table 2.**
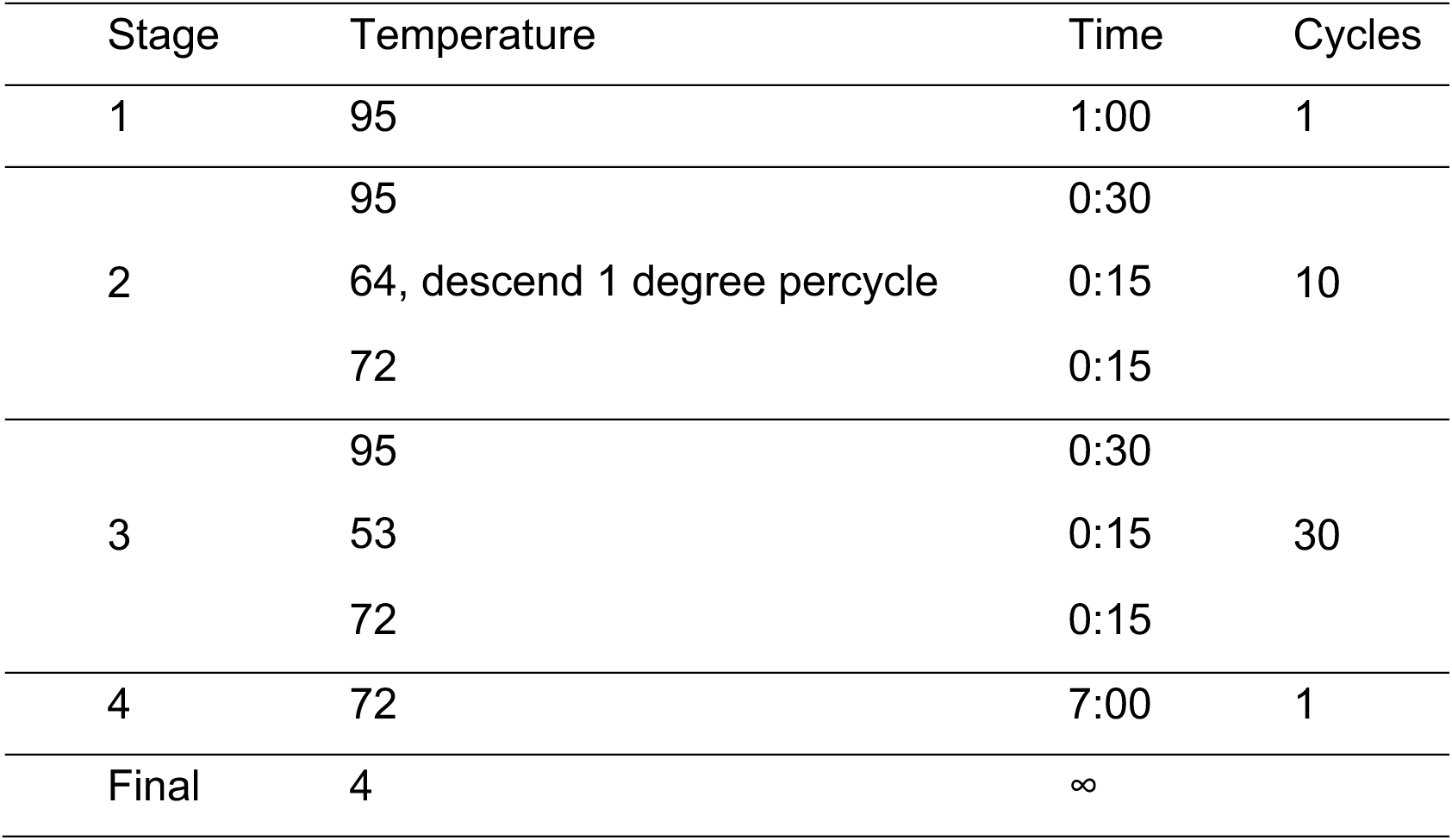
RT-PCR protocol using touchdown PCR.

**Supplementary Table 3.**
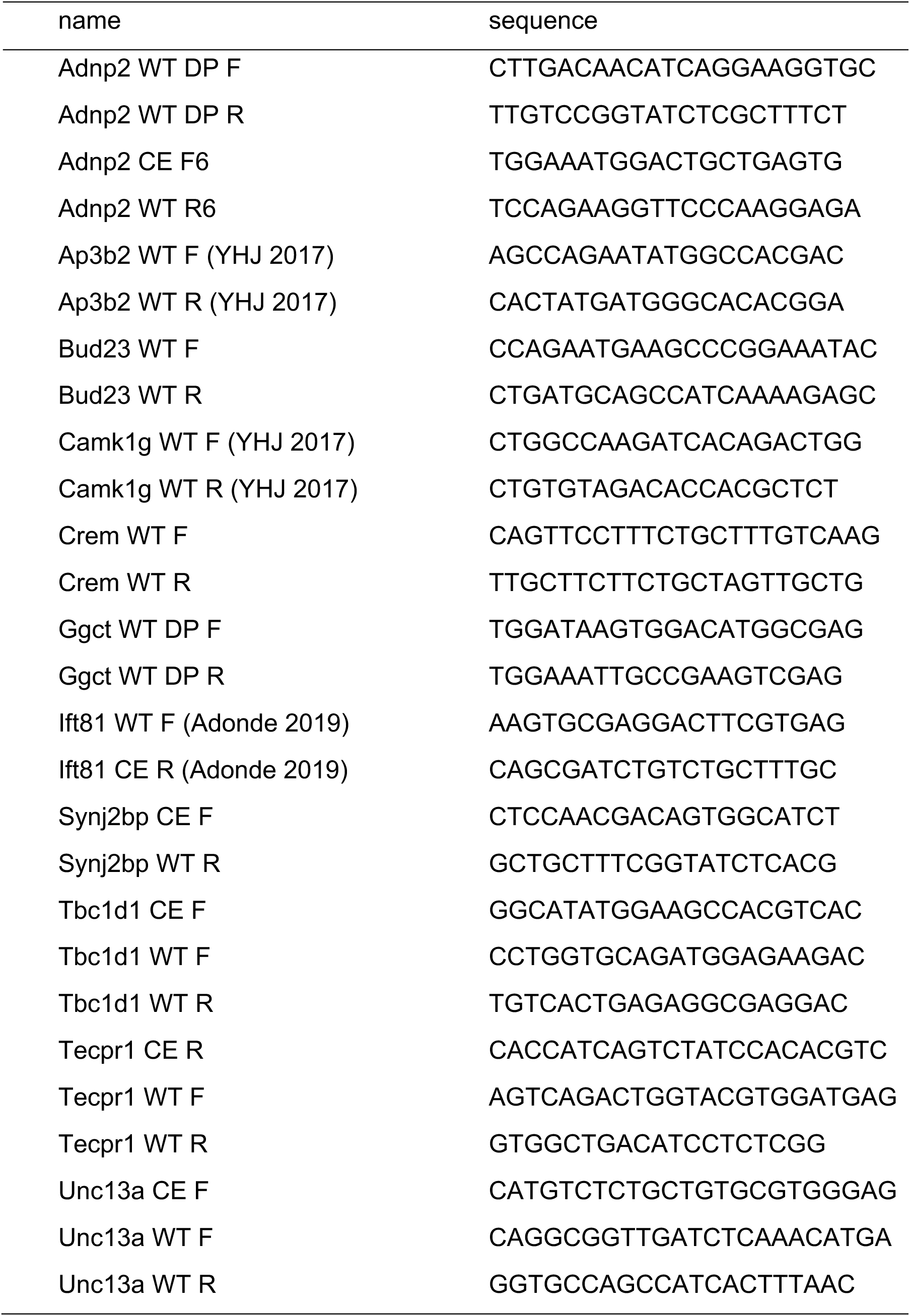
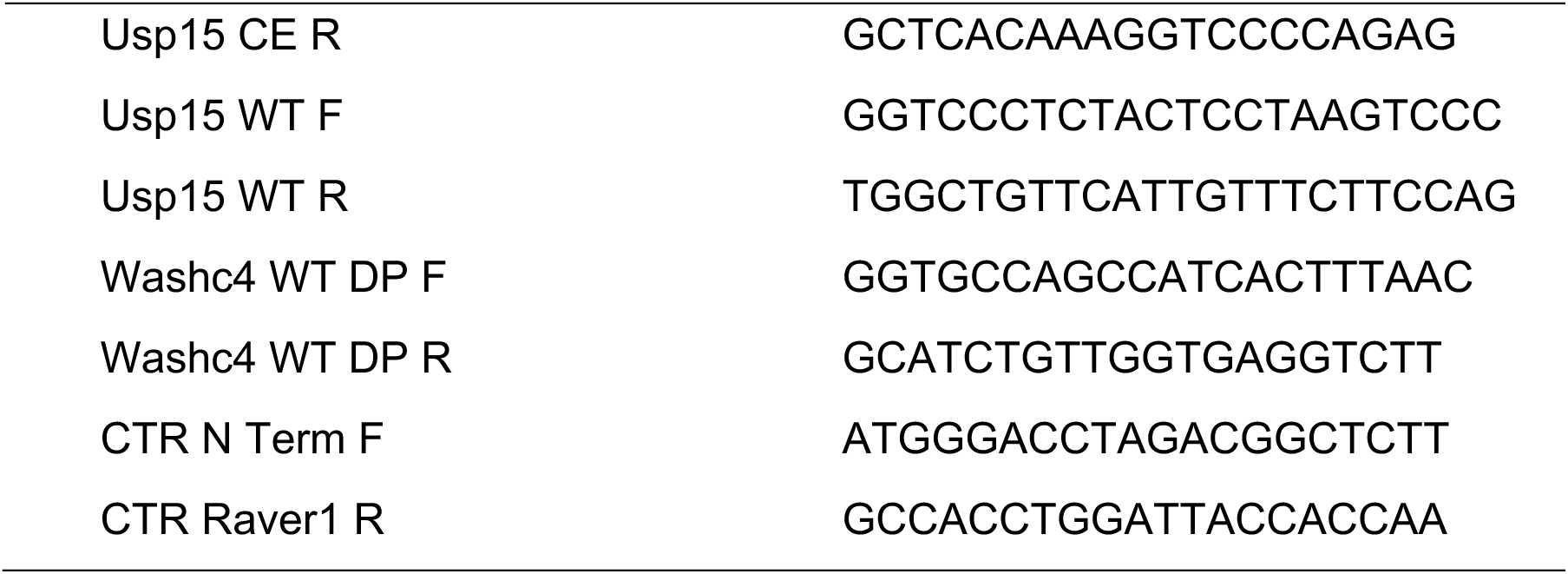
RT-PCR Primers.

**Supplementary Table 4.** Expected RT-PCR band sizes.

| Target | WT band | CE band | Single Band |
| --- | --- | --- | --- |
| Adnp2 | 187 | 339 | 128 |
| Ap3b2 | 374 | 473 | N/A |
| Bud23 | 306 | 409 | N/A |
| Camk1g | 423 | 509 | N/A |
| Crem | 332 | 476 | N/A |
| Ggct | 183 | 285 | N/A |
| Ift81 | N/A | N/A | 187 |
| Synj2bp | N/A | N/A | 326 |
| tbc1d1 | 128 | 238 | 197 |
| Tecpr1 | 329 | 382 | 358 |
| Unc13a | 169 | 213 | 166 |
| usp15 | 192 | 356 | 212 |
| Wash4c | 274 | 472 | N/A |
| CTR | N/A | N/A | 345 |

